# Drift and directional selection are the evolutionary forces driving gene expression divergence in eye and brain tissue of *Heliconius* butterflies

**DOI:** 10.1101/463174

**Authors:** Ana Catalán, Adriana Briscoe, Sebastian Höhna

## Abstract

Investigating gene expression evolution over micro- and macroevolutionary timescales will expand our understanding of the role of gene expression in adaptation and speciation. In this study, we characterized which evolutionary forces are acting on gene expression levels in eye and brain tissue of five *Heliconius* butterflies with divergence times of ~5-12 MYA. We developed and applied Brownian motion and Ornstein-Uhlenbeck models to identify genes whose expression levels are evolving through drift, stabilizing selection, or a lineage-specific shift. We find that 81% of the genes evolve under genetic drift. When testing for branch-specific shifts in gene expression, we detected 368 (16%) shift events. Genes showing a shift towards up-regulation have significantly lower gene expression variance than those genes showing a shift leading towards down-regulation. We hypothesize that directional selection is acting in shifts causing up-regulation, since transcription is costly. We further uncover through simulations that parameter estimation of Ornstein-Uhlenbeck models is biased when using small phylogenies and only becomes reliable with phylogenies having at least 50 taxa. Therefore, we developed a new statistical test based on Brownian motion to identify highly conserved genes (i.e., evolving under strong stabilizing selection), which comprised 3% of the orthoclusters. In conclusion, we found that drift is the dominant evolutionary force driving gene expression evolution in eye and brain tissue in *Heliconius*. Nevertheless, the higher proportion of genes evolving under directional than under stabilizing selection might reflect species-specific selective pressures on vision and brain necessary to fulfill species-specific requirements.

## Introduction

Species and populations diverge through the accumulation of genetic changes that affect coding or non-coding genomic regions. Genetic variation affecting gene expression has the potential of changing gene expression patterns in a spatiotemporal manner, by changing gene expression profiles in specific organs and cell types at particular developmental stages (Carroll 2005; Signor and Nuzhdin 2018). This spatiotemporal attribute of gene expression might enable evolutionary change in a compartmentalized way, allowing for change where it is required but also allowing for the needed processes to remain conserved. Phenotypic diversity caused by changes in gene expression encompasses a great variety of traits, including changes affecting an organism’s coloration (Nadeau 2016), size and shape (Ahi et al. 2017), as well as sensory perception and behavior, amongst other phenotypes (Lee et al. 2000; Wanner et al. 2007). Even though major advances have been made in linking gene expression variation to a phenotype (Catalán et al. 2016; Glaser-Schmitt and Parsch 2018), discerning the evolutionary forces shaping gene expression level variation among closely related species is an area that needs further research.

To understand the evolutionary forces acting on gene expression it is necessary to model within and between species gene expression variance. Neutral gene expression divergence between species leads to gene expression difference through divergence alone. Thus, neutral changes in gene expression modeled by random drift provides a null hypothesis to detect deviations from the expected neutral gene expression divergence. A linear relationship between divergence time and gene expression variance difference has been proposed for closely related species, assuming a clock-like (i.e., constant through time) rate of gene expression divergence (Khaitovich et al. 2004; Khaitovich 2005). Another approach to studying the evolutionary forces shaping gene expression evolution, which is motivated by statistical phylogenetics, is fitting Brownian motion (BM) models. BM-models are often used to describe changes in continuous trait through time using random drift with rate σ^2^ and taking into account the known phylogeny of the taxa of interest (Felsenstein 1985). Thus, in a BM context, σ^2^ can also be described as the volatility parameter that determines the rate at which a trait’s value diffuses away from its current state (Bedford and Hartl 2009). Fitting BM models to investigate gene expression evolution has shown that stabilizing selection and evolution through genetic drift can be readily characterized (Kalinka et al. 2010; Wong et al. 2015).

Ornstein-Uhlenbeck (OU) models have also been used to study continuous trait evolution in a phylogenetic context (Hansen 1997; Butler and King 2004). Ornstein-Uhlenbeck models, an extension to BM-models, include two extra parameters, α and θ, for modeling the strength of stabilizing (α) selection towards a phenotypic optimum (θ). As in a BM context, σ^2^ is the rate at which a trait changes through time, and α is the force pulling back the diffused trait to an optimum state. This is analogous to stabilizing selection pulling back a trait to its optimum value after having experienced a departure from it. Theta (θ) is described as the trait’s optimum state at a particular time point toward which the pull of α is aimed (Hansen 1997; Butler and King 2004). OU-models offer a useful framework to generate hypothesis about the evolutionary forces acting on transcriptome levels, whether it is drift, stabilizing selection or directional selection (Bedford and Hartl 2009; Rohlfs and Nielsen 2015; Wong et al. 2015; Chen et al. 2017; Stern and Crandall 2018).

In this study, we used five closely-related species of *Heliconius* butterflies to explore the evolutionary forces shaping gene expression variation in eye and brain tissue. *Heliconius charithonia*, *H. sara*, *H. erato*, *H. melpomene* and *H. doris* (Figure 1) belong to four of the seven distinct *Heliconius* phylogenetic clades with divergence times ranging from 5.5 to 11.8 MYA. Beside showing a great diversity in wing color patterns (Kronforst and Papa 2015), *Heliconius* butterflies also show a diversity of life history traits (Salcedo 2010; Merrill et al. 2015), mating systems (Beltrán et al. 2007; Walters et al. 2012) and behavior (Mendoza-Cuenca and Macías-Ordóñez 2005). Since *Heliconius* butterflies are diurnal species, visual stimuli provide key sources of information about the environment. For example, flowers and oviposition sites, potential mates or predators are all targets of interest to butterflies in which the first line of perception is visual (Finkbeiner et al. 2014; Finkbeiner et al. 2017). After visual cues are detected by the visual system, the detected information travels to the brain, where it is processed and its output can result in a specific behavior or physiological response. Thus, the brain’s processing and output together with the visual system have the potential of being finely tuned according to a species’ life history. In the case of *Heliconius* butterflies, a high diversity of adult compound eye retinal mosaics (between sexes and species) has been discovered (McCulloch et al. 2017), as well as species-specific differences in brain morphology (Montgomery et al. 2016). Which evolutionary forces are shaping adult eye and brain expression in *Heliconius* is one question we seek to investigate, and in that way, gain an understanding into the potential role of inter-species gene expression differences in speciation and adaptation.

**Figure 1.**
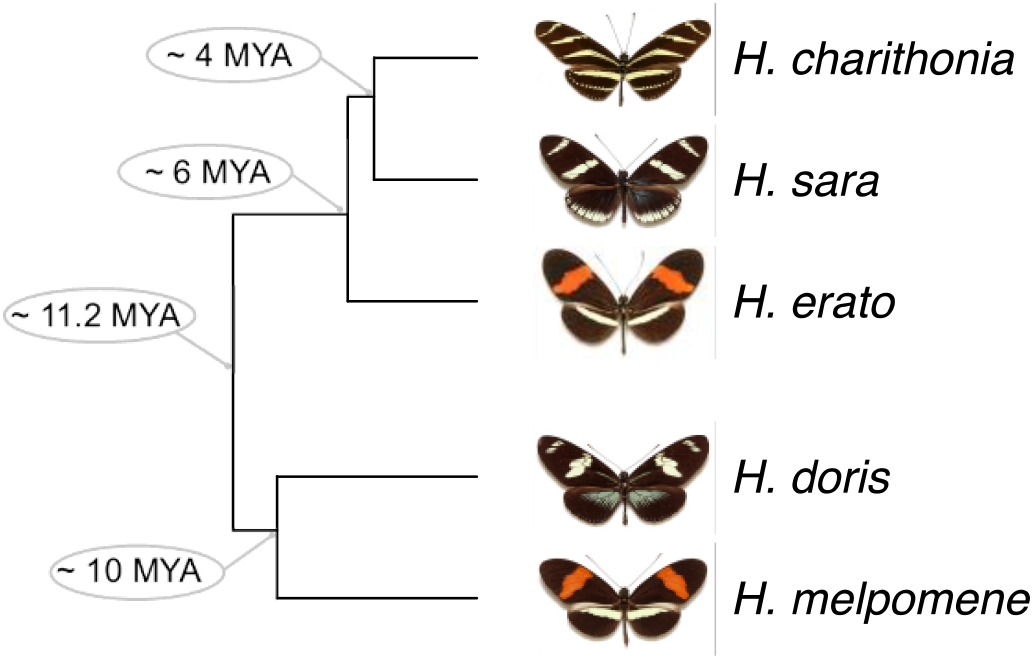
Phylogenetic relationship of the *Heliconius* species used in this study showing divergence times at each node (Kozak et al. 2015).

Therefore, in this study we investigated which evolutionary forces are driving gene expression variation in eye and brain tissue. More specifically, we aimed to identify if expression variation in individual genes is evolving, for example, through drift, stabilizing selection or directional selection. To this end, we generated a set of orthoclusters shared among our five butterfly species together with expression data for each gene in each orthocluster. We characterized the selective forces acting on gene expression levels thereby revealing the fraction of the transcriptome evolving under drift, directional selection, as well stabilizing selection.

## Methods

### Data set

The data set used in this study was published in (Catalán et al. 2018) and retrieved from ArrayExpress: E-MTAB-6810 and Dryad data identifier: DOI: doi: 10.5061/ dryad.ds21fv5. In Catalán et al (2018), whole transcriptomes were generated from eye and brain tissue combined for *Heliconius charitonia*, *H. sara*, *H. erato*, *H. doris* and *H. melpomene*. *De novo* transcriptome assemblies were generated for each species and the corresponding reads were mapped back to their matching transcriptome using Bowtie (version 1.0.0). Raw read counts and FPKM values were calculated for each species and used for downstream analysis. TransDecoder (version 5.0.2) was used to identify candidate coding regions from each *de novo* Trinity transcriptome. The predicted coding sequences were utilized to annotate each transcriptome by identifying orthologous hits in UniProt, Flybase and Pfam databases using blastp (2.2.30) and keeping only hits with an *e*-value < 10-3 (Altschul et al. 1990; Chintapalli et al. 2007; Punta et al. 2012).

### Orthology assessment

The set of orthoclusters used to assess gene expression evolution across *Heliconius* was retrieved from supplementary table S15 published in Catalán *et. al.* 2018. Briefly, the Unrooted Phylogenetic Orthology (UPhO) pipeline and model was used to assess orthologous relationships between the five *Heliconius* species (Ballesteros and Hormiga 2016). UPhO uses an all species pairwise blastp search and a Markov clustering algorithm (MCL) (version 1.0.0) (Enright et al. 2002) to cluster sequences according to sequence similarity. Clustered sequences were aligned with MAFFT (version 7.3.05) (Katoh and Toh 2008) and curated after alignment with trimAl (version 1.3) (Capella-Gutiérrez et al. 2009). Phylogenetic inference for each sequence cluster was done using RAxML (version 8.2.10) (Stamatakis 2006) and orthology was assessed for each generated tree using the UPhO algorithm. A matrix with log_2_ FPKM values was generated for each orthoclusters which was used to analyze gene expression variance.

### Modelling Gene Expression Evolution

To study the forces driving gene expression evolution, we implemented a set of six different statistical models (Table 1). Each model models the mean species gene expression level (between-species variance) and the gene expression levels of individual samples per species (within-species variance). How these mean species gene expression levels evolve, or not, along the phylogeny and over time, is specific and central to each model. We estimated the parameters of each model and performed Bayesian model selection using Bayes factors to establish which model describes the observed data best and thus which process is most likely to drive gene expression evolution in the five *Heliconius* species of our study (see below).

**Table 1.** Summary of the models implemented in this work

| Model | Description | Parameters |
| --- | --- | --- |
| **Equal species means** | All species have the same mean gene expression level. | $\mu$ – global mean gene expression level |
| **Unequal species means** | All species have their own independent mean gene expression level. | $\mu_i$ – mean gene expression level per species |
| **Brownian motion (BM)** | Random drift of the species mean gene expression level along the phylogeny. | $\sigma^2$ - rate of drift |
| **Brownian motion (BM) with shift** | Random drift with one branch having a different rate (directional selection). | $\sigma_B^2$ - rate of drift background branch |
| | | $\sigma_F^2$ - rate of drift foreground branch |
| **Ornstein-Uhlenbeck (OU)** | Stabilizing selection of the species mean gene expression level evolving along the phylogeny. | $\sigma^2$ - rate of drift<br>$\alpha$ - strength of selection<br>$\theta$ - optimal gene expression level |
| **Ornstein-Uhlenbeck (OU) with shift** | Directional selection due to a shift in optimal gene expression level. | $\sigma_B^2$ - rate of drift background branch<br>$\sigma_F^2$ - rate of drift foreground branch.<br>$\theta_B$ - optimal gene expression level background branch<br>$\theta_F$ - optimal gene expression level foreground branch<br>$\alpha$ - strength of selection |

The simplest model of gene expression assumes that all species have the exact same mean gene expression level. In this case, we only have one parameter μ which defines the mean gene expression level of all species. The expression level X_ij_ of individual *i* from species *j* is modeled using a normal distribution with X_ij_ ~ Norm(μ, δ^2^_i_). We chose a uniform prior distribution between −20 and 20 for the mean gene expression parameter μ. Note that we assume that every species has its own gene expression variance parameter δ^2^_i_ (see below). This model assumes there is no evolution of gene expression levels, i.e., gene expression levels are completely conserved among species.

The second model that we implemented was a model where each species has its own gene expression mean μ_i_. Thus, we model the gene expression level X_ij_ of gene *i* from species *j* using a normal distribution with X_ij_ ~ Norm(μ_i_, δ^2^_i_). In this model, each species has a different mean gene expression level, but these gene expression levels do not evolve under an evolutionary model; they are intrinsically different without any mechanistic reason (no phylogenetic signal). As with the first model, we assumed a uniform prior distribution between −20 and 20 for each mean gene expression level μ.

The third model that we implemented was a phylogenetic Brownian motion model (Felsenstein 1985). We assume that any gene expression value at the root of the phylogeny is equally probable. Then, the mean gene expression levels μ evolve along the lineages of the phylogeny. The Brownian motion model specifies that the focal variable, μ in our case, is drawn from a normal distribution centered around the value of the ancestor, μ_A_. The amount of change, i.e., rate of random drift, is defined by the parameter σ^2^. We assumed a log-uniform prior distribution between 10E-5 and 10E5 for the drift parameter σ^2^. Thus, the mean gene expression levels μi for the species of the phylogeny are distributed according to a multivariate normal distribution where the covariance structure is defined by the phylogeny (Felsenstein 1985). This means that more closely-related species are expected to have a more similar mean gene expression level because they share more evolutionary history (i.e., they are more recently diverged), which is modeled by the covariance structure. Such a phylogenetic model of gene expression evolution has been applied previously by (Bedford and Hartl 2009). Importantly, the BM model only defines how the mean gene expression levels evolve but does not allow for any sample variance of the individuals of a species. Therefore, we extended the standard phylogenetic BM model to allow for within-species sample variance where again the expression level X_ij_ of individual *i* from species *j* is normally distributed with X_ij_ ~ Norm(μi, δ^2^_i_) where δ^2^_i_ is the within-species variance parameter (Ives et al. 2007; Rohlfs and Nielsen 2015). This extension to allow for within-species variance was developed for all phylogenetic models (BM, BM with shift, OU and OU with shift).

The fourth model that we implemented was a phylogenetic BM model with branch-specific rates of evolution, thus detecting directional selection. The mean gene expression level evolves under a BM model (i.e., random drift) where the rate of evolution for branch *k* is given by σ^2^_k_ (O’Meara et al. 2006; Eastman et al. 2011). Thus, a branch with a higher rate of evolution σ^2^_k_ signifies more change in gene expression levels than under a constant rate random drift model (the BM model). Directional selection can therefore be detected by inferring an elevated estimate of σ^2^_k_ compared with the background rate of drift σ^2^. Specifically, we applied a background rate of drift σ^2^_B_ to all branches except the chosen foreground branch which received its own rate of drift σ^2^_F_.

The fifth model we implemented was a phylogenetic Ornstein-Uhlenbeck process (Hansen 1997). The Ornstein-Uhlenbeck (OU) process models, similar to BM, the evolution of the mean gene expression level per species along a phylogenetic tree. However, unlike BM, the mean expression level diffuses with rate σ^2^ and is attracted with strength α to an optimum level θ. Thus, the OU process has an expected variance of σ^2^/2α which is independent of time, i.e., does not increase with increasing time but instead stabilizes (c.f. Figure 10). The variance becomes small if either the strength of selection is large or the rate of drift is small. This is, in fact, a major problem for OU models which cannot distinguish if attraction (or selection) is strong or diffusion is weak (Ho and Ané 2014; Cooper et al. 2016).

The sixth model we implemented was an Ornstein-Uhlenbeck process with a branch-specific shift in both the rate of drift σ^2^ and the optimum gene expression level θ (Rohlfs et al. 2014; Uyeda and Harmon 2014). Thus, this branch-specific OU model is analogous to the branch-specific BM model, allowing for directional selection in an OU framework. Specifically, we tested if there was a significant support for the chosen foreground branch which received its own rate of drift σ^2^_F_ and optimum θ_B_ to be different from the background rate of drift σ^2^_B_ and optimum θ_B_. We used the same prior distributions as before and assumed that both parameters for the background and foreground branches are drawn from the identical prior distribution. This model has in total five free additional parameters along with the five nuisance parameters (the within-species variances). Thus, we expect that this model is more prone to be over-parameterized for our dataset with five species. Nevertheless, our Bayesian approach for parameter estimation and model selection integrates over parameter uncertainty and penalizes extra parameters by integrating over the prior distribution.

### Parameter Estimation and Model Selection

We estimated parameters for our different models in a Bayesian statistical framework. Thus, we approximated the posterior distribution of the model parameters using Markov chain Monte Carlo sampling (Metropolis et al. 1953; Hastings 1970). We ran a separate MCMC analysis for each model and gene, 2393 analysis per model. Every model parameter was updated twice per MCMC iteration where the order of parameter updates was chosen randomly. We applied the same settings of the MCMC algorithm for each model. First, a burn-in phase of the MCMC algorithm was run for 2,000 iterations with auto-tuning every 100 iterations. Then, the actual MCMC simulation was run for 50,000 iterations with sampling 10 iterations, yielding 5,000 samples from the posterior distribution (Höhna et al. 2017).

Model selection was performed using marginal likelihoods. Marginal likelihoods are the probability of the data for a specific model integrated over all possible parameter values. From the marginal likelihood we can then compute Bayes factors and model probabilities (i.e., weights of a model being the true model generating the data given a set of candidate models). We approximated the marginal likelihoods using stepping stone sampling (Fan et al. 2011). The stepping stone algorithm implemented in RevBayes consisted of 128 MCMC runs where each MCMC ran had the likelihood function raised to the power of β computed by the quantiles of a beta probability distribution (Höhna et al. 2017).

### Data availability

The five different models that we used in our study are implemented in Bayesian phylogenetic inference software RevBayes v1.0.8 (Höhna et al. 2016). For efficient computations, we implemented the restricted maximum likelihood (REML) algorithm for BM models (Felsenstein 1985) and OU models (Fitzjohn 2012; Freckleton 2012). The source code and compiled executables of RevBayes are available from https://github.com/revbayes/revbayes and tutorials about the analyses are available from https://revbayes.github.io/tutorials/.

## Results

To assess the evolutionary forces acting on gene expression levels in *H. charithonia*, *H. sara*, *H. erato*, *H. doris* and *H. melpomene* (Figure 1), we used the gene orthology dataset, composed of 2373 orthologous genes, published before (Catalán et al. 2018). From this previous work, we also obtained FPKM (<u>F</u>ragments <u>p</u>er <u>K</u>ilobase per <u>M</u>illion mapped read) values for each gene and sample. The log_2_ transformed FPKM values were used to build an expression matrix and to model gene expression evolution.

### Testing for equality of within-species variance in gene expression levels

Equality of variances among populations or samples drawn from a normal distribution is usually assumed when testing for differences in mean values obtain from continuous data or gene expression data as in our case (Warnefors and Eyre-Walker 2012; Rohlfs et al. 2014). Assuming equality of variances when it is not the case can lead to high Type I error rates (Gastwirth et al. 2009). To avoid assuming equality of variances, we computed the within-species variances for each orthocluster and checked for the presence of a correlation across species. From a pairwise assessment of within-species variance we found no significant correlation among all possible pairs, with Pearsons’s rho values ranging from 0.07 to 0.2 (Figure 2). Since gene expression variance across species is heterogeneous, hence not correlated among species, we treated within-species variance as a random variable when fitting BM and OU models.

**Figure 2.**
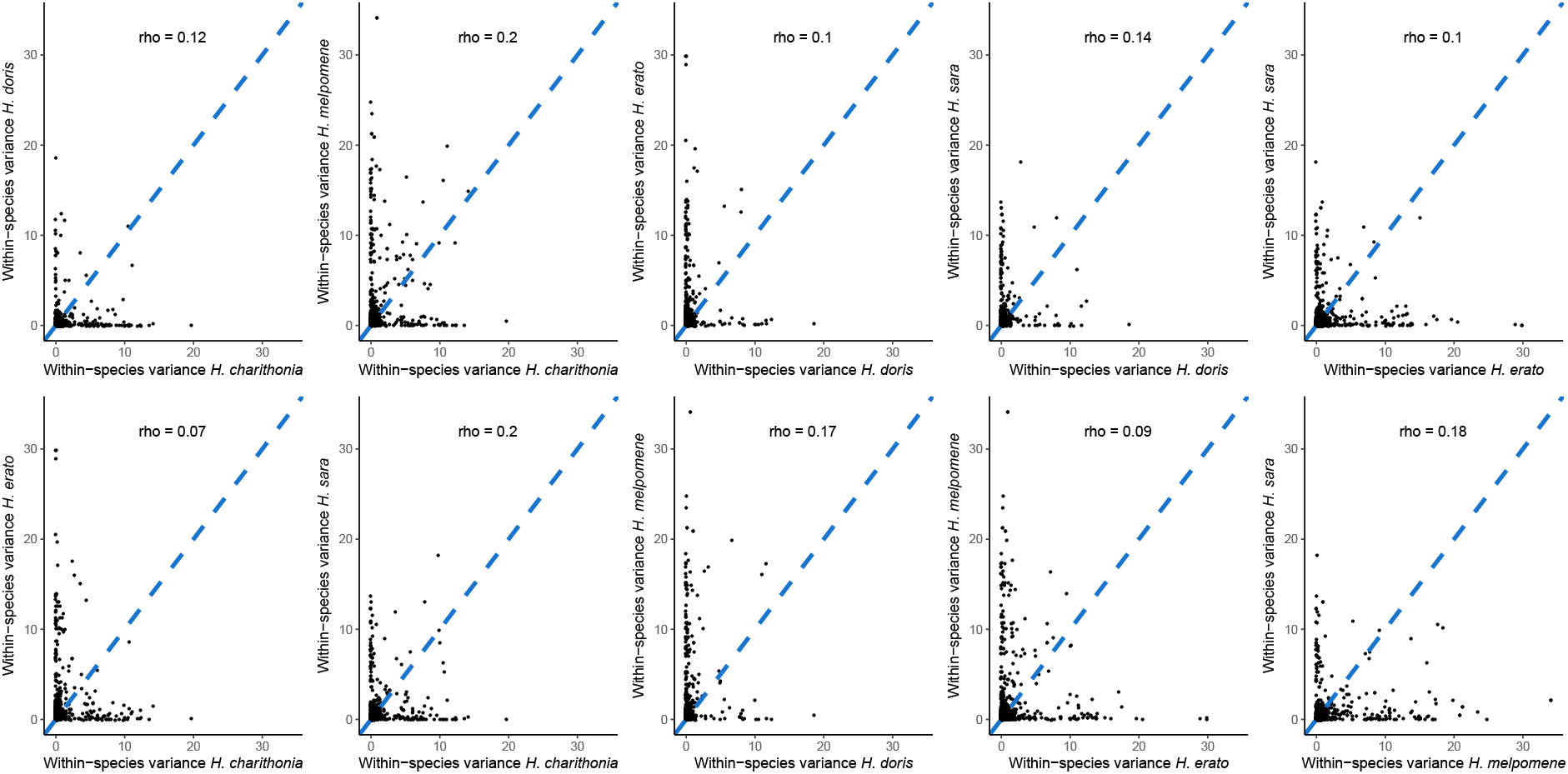
Pairwise correlation between five *Heliconius* species and their gene expression within-species variance. The correlation strength between within-species variances was estimated by calculating Pearson’s correlation coefficient, rho ranging from 0.07 to 0.2.

### Testing for gene expression evolution through drift using Brownian motion models

We applied BM models to describe changes in gene expression levels through random drift. As the alternative hypotheses, we used two non-phylogenetic models where either all species had identical mean gene expression levels (Model 1) or all species had their own independent mean gene expression levels (Model 2). For each gene we computed the probability of the BM model having produced the observed data, i.e., a high probability means that it is more probable that the gene expression levels evolved under a BM model whereas a low probability means that it is more probable that the gene expression levels evolved under a non-phylogenetic model (Model 1 and Model 2). A model probability of >0.75 corresponds to a Bayes factor of >3 (positive support) and a model probability of >0.95 corresponds to a Bayes factor of >20 (strong support).

Our results show that the majority of gene expression levels (2369 out of 2393) are evolving as expected given the known *Heliconius* species phylogeny, i.e., that there is a strong phylogenetic signal (Figure 3). These results indicate that drift is the dominating evolutionary force driving transcriptome change. Only a small fraction of the orthoclusters did not show a phylogenetic signal, opening the question of the putative genetic forces shaping gene expression levels of these genes.

**Figure 3.**
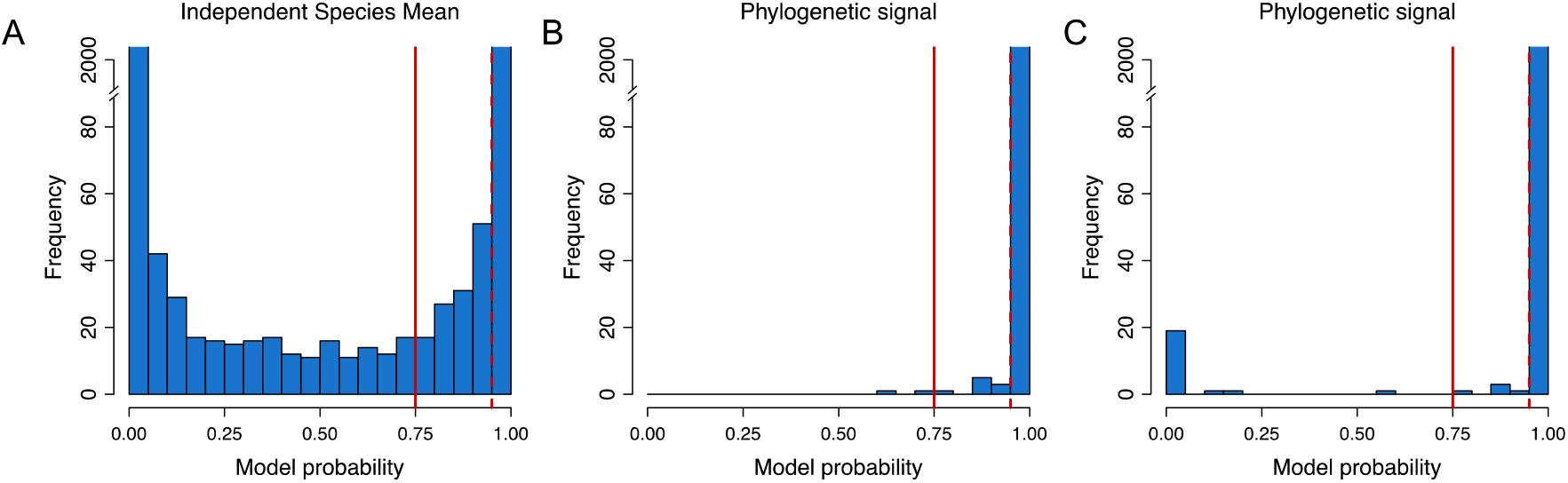
Testing for random drift in gene expression levels of *Heliconius* using Brownian motion. Significance is shown at model probability > 0.75 (solid red, Bayes factor > 3, positive support) and model probability > 0.95 (solid red, Bayes factor > 20, strong support). (A) Shows the comparison between the two non-phylogenetic models (identical vs independent species mean). (B) Shows the model probability of the BM model compared with the independent species mean model. (C) Shows the model probability of the BM model compared with the identical species mean model.

### Testing for conserved gene expression level

The next question we explored was how prevalent conserved gene expression levels are in eye and brain tissue of *Heliconius*. This question could be answered with our previous results by computing how often Model 1, with identical species means, was recovered (Figure 2C). However, our model selection procedure relied on computing marginal likelihoods which are intrinsically sensitive to the choice of prior distribution (Berger 1990; Kass and Raftery 1995; Sinharay and Stern 2002). Therefore, we additionally performed a sensitivity analysis of σ^2^ = 0 using Monte Carlo simulation as follows (Goldman 1993). We estimated the posterior distribution of all parameters under the identical species mean model (the only parameters were the within-species variances), then we used 1,000 parameter samples from the posterior distribution to simulate gene expression datasets (e.g., a dataset consisting of a single gene with five species and 6-12 individuals per species) under the identical species mean model, yielding 1,000 simulated datasets per gene in total. Then, for every gene of the 2393 genes, we estimated σ^2^ for each simulated dataset as well as the original dataset, which amounted to a total of 2,395,393 MCMC analyses. Finally, we calculated if the mean posterior estimate of the empirical dataset was larger than 95% of the mean posterior estimates of the simulated datasets. In the cases when the mean posterior estimate of σ^2^ was not larger than the mean estimate of 95% of the simulated datasets we concluded that these genes are highly conserved (Figure 4).

**Figure 4.**
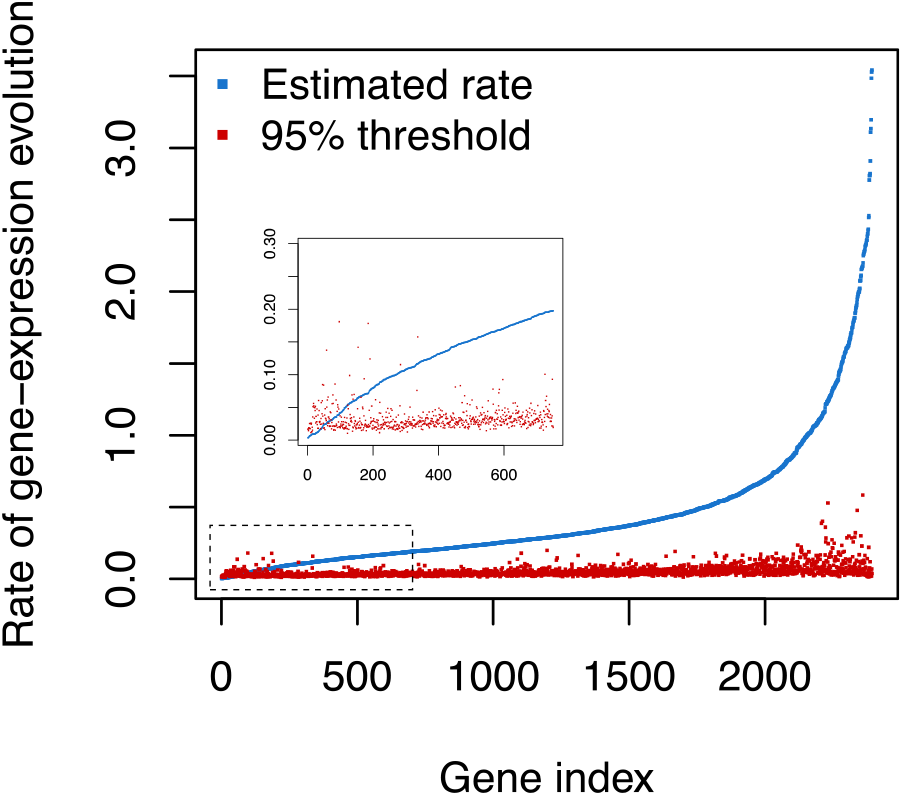
Posterior mean estimates of the drift rate σ^2^ (blue) and the 95% threshold computed (red) using Monte Carlo simulations. The genes were sorted by an ascending estimate of σ^2^. Inset: close-up of genes whose σ^2^ is not significantly bigger than zero.

By using the described approach, we uncover a set of 83 orthoclusters whose gene expression variance across species is highly conserved (Figure 4 and Figure S1). A sigma squared not significantly different from zero, can be caused by stabilizing selection hindering gene expression divergence, resulting in more similar gene expression patterns across different *Heliconius* species.

### Testing for stabilizing selection acting on gene expression levels

Subsequently, we moved forward into implementing an Ornstein-Uhlenbeck model (OU) to investigate the strength of stabilizing selection (Bedford and Hartl 2009; Rohlfs et al. 2014). OU-models are an extension of BM-models, in which they include two extra parameters, α and θ. In a BM context, if σ^2^ is the rate at which a trait changes through time, α is then described as a force pulling back the diffused trait to an optimum state (θ).

We estimated the marginal likelihood for each gene under a BM model and an OU model. Then, we computed the probability (i.e., support) of an OU model over a BM model using the marginal likelihoods. Our results show a very low support for stabilizing selection (Figure 5). When the marginal likelihoods were examined, in 99.7% of the cases a BM model explained our expression data better than an OU-model.

**Figure 5.**
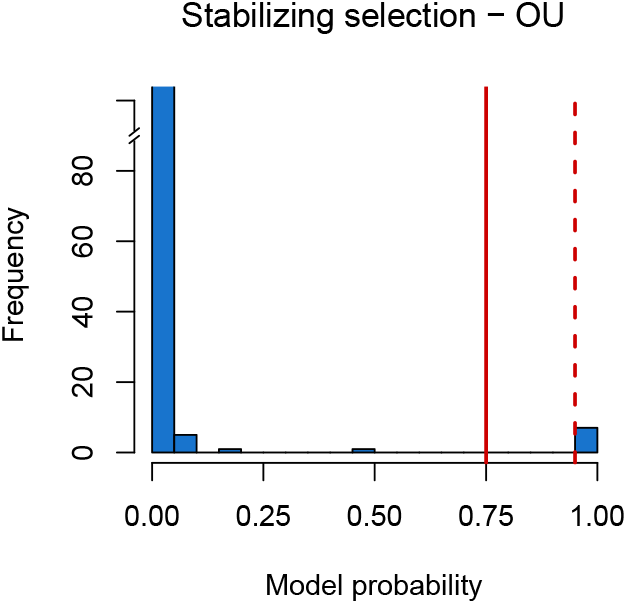
Model probability when testing the strength of alpha when fitting an OU model for the assessment of stabilizing selection. Significance is shown at model probability > 0.75 (solid red, Bayes factor > 3, positive support) and model probability > 0.95 (solid red, Bayes factor > 20, strong support). There are only 7 genes with a significant support for stabilizing selection.

### Testing the power to estimate stabilizing selection

Our results indicating that very few genes evolved under stabilizing selection conflict with previous findings (Bedford and Hartl 2009). However, it has previously been discussed that when working with small phylogenies (less than ten species) there is a lack of power for parameter estimation when using an OU-model (Rohlfs et al. 2014). By simulating data under an OU model using phylogenies with varying numbers of taxa, we were able to show how parameter estimation is biased. The attraction parameter α could only be estimated closely to the true values used for the simulations when the phylogenies contained 50 or more taxa. Thus, we can observe that the bias observed for parameter estimation drops considerably when the number of taxa composing the phylogeny reaches 50 (Figure 6). This observation holds as well for the estimation of σ_2_ under a range of sigma values (Figure S2). Our simulation study shows that attention needs be paid when applying OU models to assess gene expression evolution for phylogenies containing less than 50 taxa.

**Figure 6.**
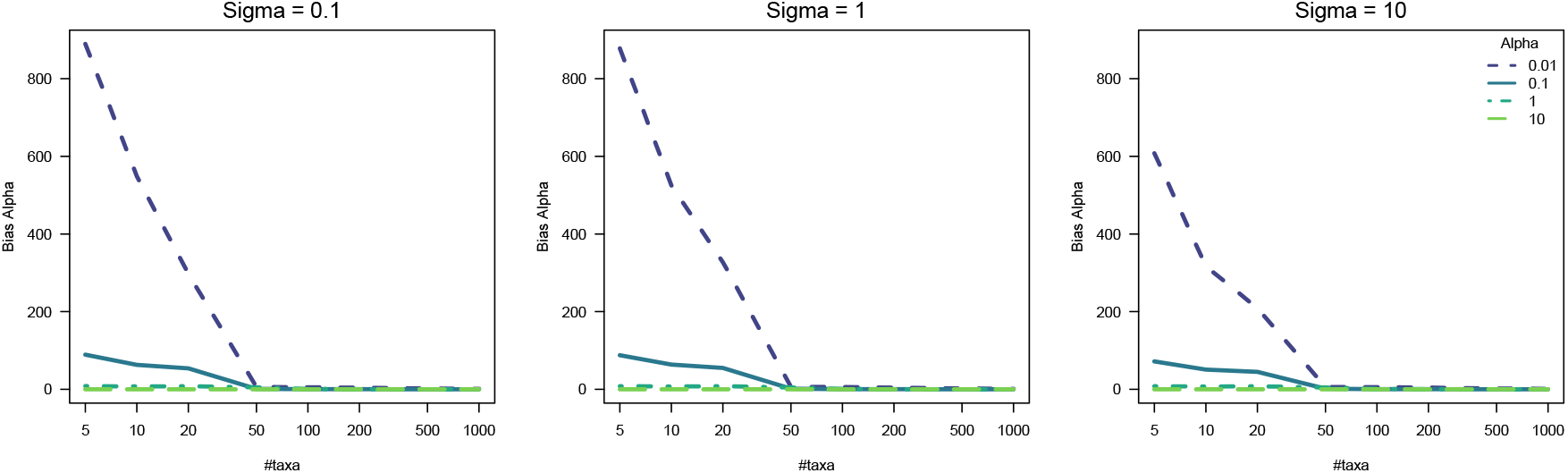
Simulation study for the assessment of parameter estimation bias under an OU-model. The relative bias in estimates of the attraction/selection parameter (α) through 1000 simulations under sigma values ranging from 0.1 – 10 and alpha values ranging from 0.01 – 10. Simulations were performed for phylogenies with sizes ranging from 5 to 1000 taxa.

### Detection of branch specific shifts in gene expression

To reveal genes whose gene expression patterns have putatively been shaped by directional selection, we tested for branch-specific shifts in evolutionary rates along the *Heliconius* tree. To explore branch-specific shifts in gene expression, firstly we used a BM model to test for the evolutionary rate (σ^2^) of a focal branch being different from the background rate (i.e., the rest of the branches in the phylogenetic tree) and assessed significance by applying Bayes factors (Figure S3 and Figure 7). Secondly, we also tested branch-specific shifts through an OU model and tested for a branch-specific shift in gene expression level optimum (θ_F_) vs the rest of the tree’s θ_B_ (Figure S4).

**Figure 7.**
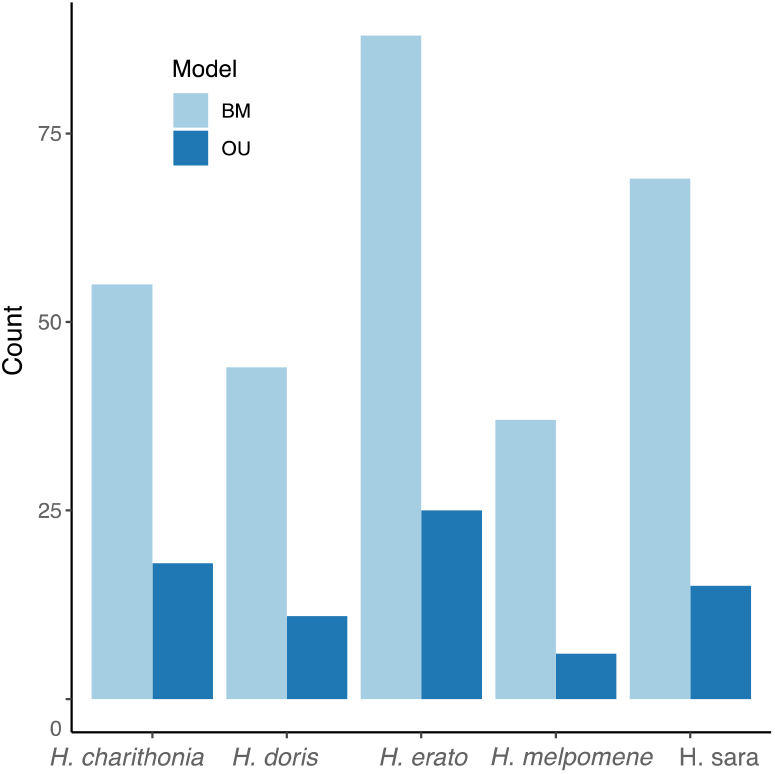
Barplot showing branch-specific shifts on gene expression levels in *Heliconius.* Bars in light blue show branch shifts identified by BM and dark blue bars show branch-shifts identified by OU models.

With a BM approach, we were able to detect a total of 322 branch specific shifts, when considering only tip branches (Figure 7). We found 112 branch specific shifts in the HER linage, 70 in HAS, 67 in HCH, 44 in HDO and 29 in HME (Figure 7 and Figure S4). HCH, HAS and HDO had more shifts towards a down-regulation, although only in HCH and HAS was this difference significant (sign test, HCH: *P*-value 6.738e-05 and HAS: *P*-value 1.653e-06). In HER and HME more up-regulated genes were causing a branch specific shift, although no significant difference was found.

When implementing an OU model we recover a total of 75 genes showing a branch specific shift in gene expression optimum (Figure 7 and Figure S4). From these genes, 55 also show a branch specific shift when implementing a BM model and 20 genes show uniquely a gene expression level shift in optimum when using an OU model (Figure S4).

Next, we assessed within-species gene expression variance of all the genes identified as having a branch-specific shift in gene expression through BM and OU models. When we plotted the distribution of the within-species variance we found that up-regulated genes have a significantly lower variance when compared to genes with a gene expression shift towards a down-regulation (Figure 8). Different evolutionary forces acting on shifts causing up- or down-regulation have the potential of maintaining different levels of gene expression variation within species.

**Figure 8.**
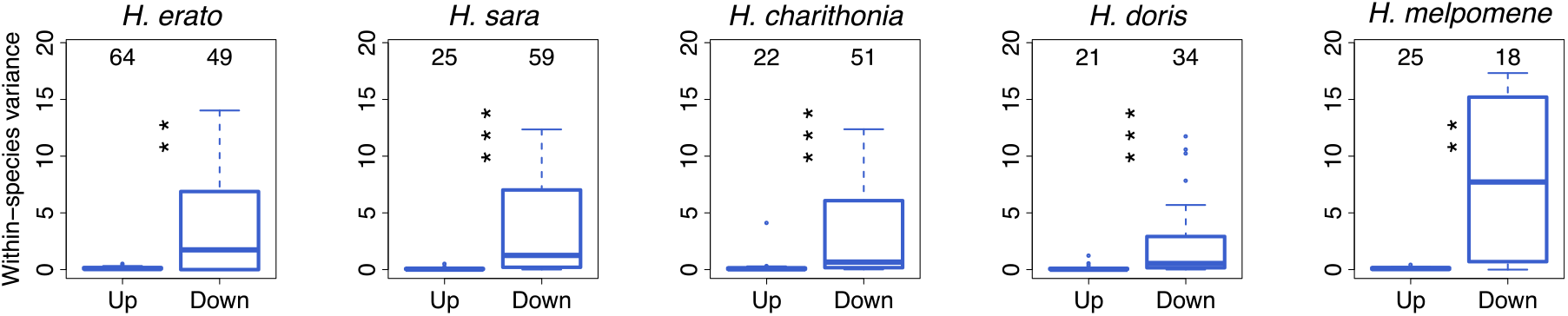
Boxplot showing the distribution of the within-species variance of gene expression levels identified as having a shift towards up-regulation and shifts towards down-regulation. Numbers above the boxplots show the total number of genes identified with a BM and an OU model. Wilcoxon-test: * *P*<0.05, ** *P*<0.01, *** *P*<0.001.

## Discussion

### Gene expression evolution through genetic drift

Our study of the evolutionary forces acting on gene expression in eye and brain tissue of *Heliconius* butterflies revealed that most transcriptome levels (81%) are evolving under drift. According to neutral expectations, phenotypic changes are expected to accumulate as a function of time, by drift and mutation alone (Lande 1976). As a consequence, the change of transcriptomic levels through drift should reflect the divergence history of the taxa of interest. From our BM analysis, we conclude that in most of the gene expression levels on eye and brain a phylogenetic signal can be recovered (Figure 3).

Consequently, we hypothesize that gene expression variation influencing phenotypic variation across species mainly arises through random drift. Evolutionary rates of gene expression evolution have been investigated at the population and at the species level and it has been found that the proportion of the type of evolutionary force acting on transcriptomic levels is not constant across taxa (Whitehead and Crawford 2006a; Nourmohammad et al. 2017; Stern and Crandall 2018). For example, when examining the evolutionary forces acting on gene expression levels in several fish populations, the authors reported that the dominant force driving expression changes was genetic drift (Whitehead and Crawford 2006b). Comparably, in a study concerning primates, genetic drift was the main force driving gene expression evolution (Khaitovich et al. 2005). The proportion of gene expression levels evolving by drift depends on the strength of natural selection acting on the interrogated transcriptome. For example, in a comparison between different organ types in mammals, gonad gene expression showed the lowest phylogenetic signal when compared to other organs like cerebellum or heart (Brawand et al. 2011). In *Heliconius* butterflies, other organs would need to be tested in order to get a more global understanding on how gene expression is evolving in the whole organism.

We explored the gene expression data further by comparing the expected gene expression divergence under a BM model to the observed data. Consequently, we simulated expression levels for 10,000 genes along the known *Heliconius* phylogeny and computed the mean of the pairwise species difference. Similarly, we computed the mean pairwise difference of the observed gene expression data. Alternatively, we can also derive the expected divergence in gene expression levels between two species over time under Brownian motion. Both species evolve under random drift and, thus, their gene expression values are normally distributed with variance σ^2^ x *T* where *T* is the time since the most recent common ancestor of the two species. Therefore, the difference in gene expression levels between the two species is normally distributed with variance 2 x σ^2^ x *T*. Since we are only interested in the absolute value of the gene expression difference, we use a truncated normal distribution instead. From this truncated normal distribution with mean zero and variance 2 x σ^2^ x *T* we compute the expected gene expression difference through time (Figure 9). For the empirical data, we estimate σ^2^ using a sum of squares approach. We find that our observed gene expression data has a close fit to the simulated data (Figure 9).

**Figure 9.**
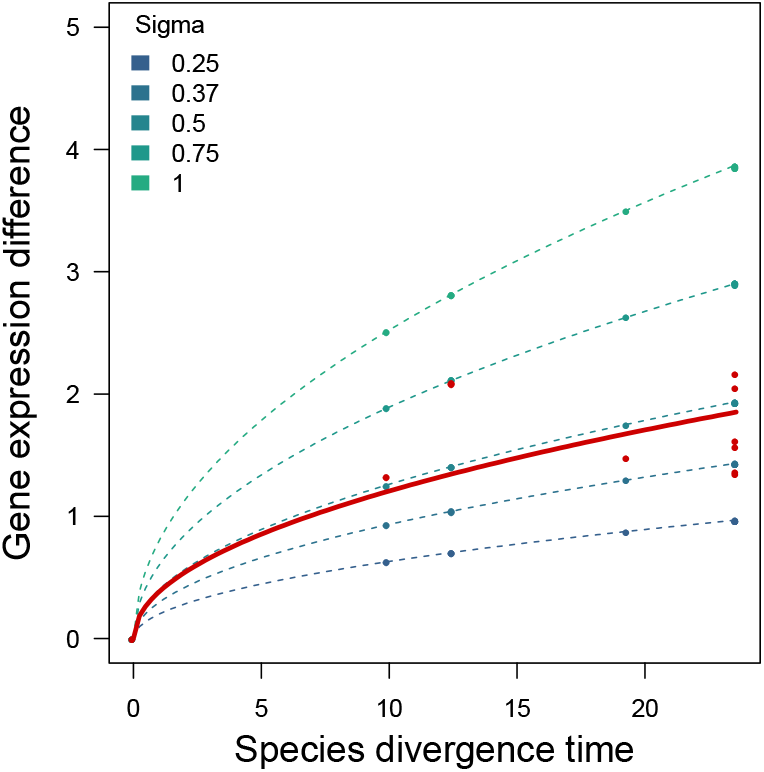
Between species gene expression variance plotted as a function of divergence time according to the *Heliconius* phylogeny. Red: σ^2^ from gene expression levels observed in *Heliconius*. Blue: simulated gene expression difference under random drift with different values of sigma.

### Gene expression evolution through genetic stabilizing selection

Studies done in *Drosophila* and mammals have shown that stabilizing selection is the evolutionary force dominating gene expression evolution (Rifkin et al. 2003; Rohlfs and Nielsen 2015). In contrast to these studies, in *Heliconius* we uncovered that only 3% of gene expression are either highly conserved (Figure 4) or evolving through stabilizing selection (Figure 5). Factors like tissue type, gene functionality turnover or epistatic levels, have the potential to influence the degree of stabilizing selection acting on the transcriptome (Larracuente et al. 2008; Kalinka et al. 2010; Romero et al. 2012). Additionally, in groups that have experienced an adaptive radiation, like in *Heliconius* (Kozak et al. 2015), and thus have recently experienced an elevated rate of trait evolution, directional selection might be more recurrent than stabilizing selection.

OU models are suitable models to study the force of stabilizing selection acting on a phenotype since the α parameter simulates the strength of selection keeping a trait close to an optimum (Beaulieu et al. 2012), as several studies exemplify (Kalinka et al. 2010; Brawand et al. 2011; Stern and Crandall 2018). When we applied an OU model to identify stabilizing selection on gene expression, we detected parameter estimation biases as shown by our simulation study (Figure 6). For small phylogenies, accurate parameter estimation is challenging since statistical power is weak with small sample sizes (Rohlfs et al. 2014). Specifically, it is very challenging with small phylogenies to distinguish between conserved gene expression levels due to low values of drift (i.e., no change) and high values of selection (i.e., drift is removed due to selection). Not only the number of taxa, but also the depth of the phylogeny can influence the suitability of OU-models to infer stabilizing selection (Fay and Wittkopp 2008; Bedford and Hartl 2009). Therefore, we propose that for small phylogenies, testing for σ^2^ =0 under a BM framework and assessing for significance by applying Monte Carlo simulations is a better model choice to uncover stabilizing selection. When using this approach, we identified 83 genes with conserved gene expression levels across species. These genes might be involved in maintaining conserved processes that are essential for the function of eye and brain tissue in *Heliconius*. For example, from the top ten genes with the most conserved gene expression levels, we found the transcription factor *bobby sox* (*bbx*) (Group_674, Appendix 1). BBX, belongs to the high mobility box domain superfamily, which are involved in transcription, replication and chromatin remodeling (Chintapalli et al. 2007). BBX has also been found to have orthologues in flies, human and mice (Nitta et al. 2015) suggesting a high essentiality of *bbx* expression. Another highly conserved orthocluster (Group 977, Appendix 1) was annotated as *glaikit* (Chintapalli et al. 2007), which is known to be essential for the formation of epithelial polarity and nervous system development (Dunlop et al. 2004).

We repeated our previous data exploration by simulating 10,000 genes under an OU-model under a range of σ^2^ and α values and computed the mean differences between pairs of species. We observed a reasonably good fit to our data (Figure 10). Most importantly, in Figure 10 we observed a steeper change in gene expression difference between closely related species (low evolutionary distance). Consequently, adding more species, including closely related species, to our model could not only improve OU parameter estimation but could also help in disentangling the evolutionary forces acting on gene expression divergence, specially between closely related species. Interestingly, the observed differences (red dots) in gene expression levels (which are averaged over all 2393 genes) show a clear departure from the expected difference predicted by both a BM and OU model (red lines in Figure 9 and Figure 10). This could be a strong indication of gene expression divergence under different evolutionary forces (drift, stabilizing selection or directional selection).

**Figure 10.**
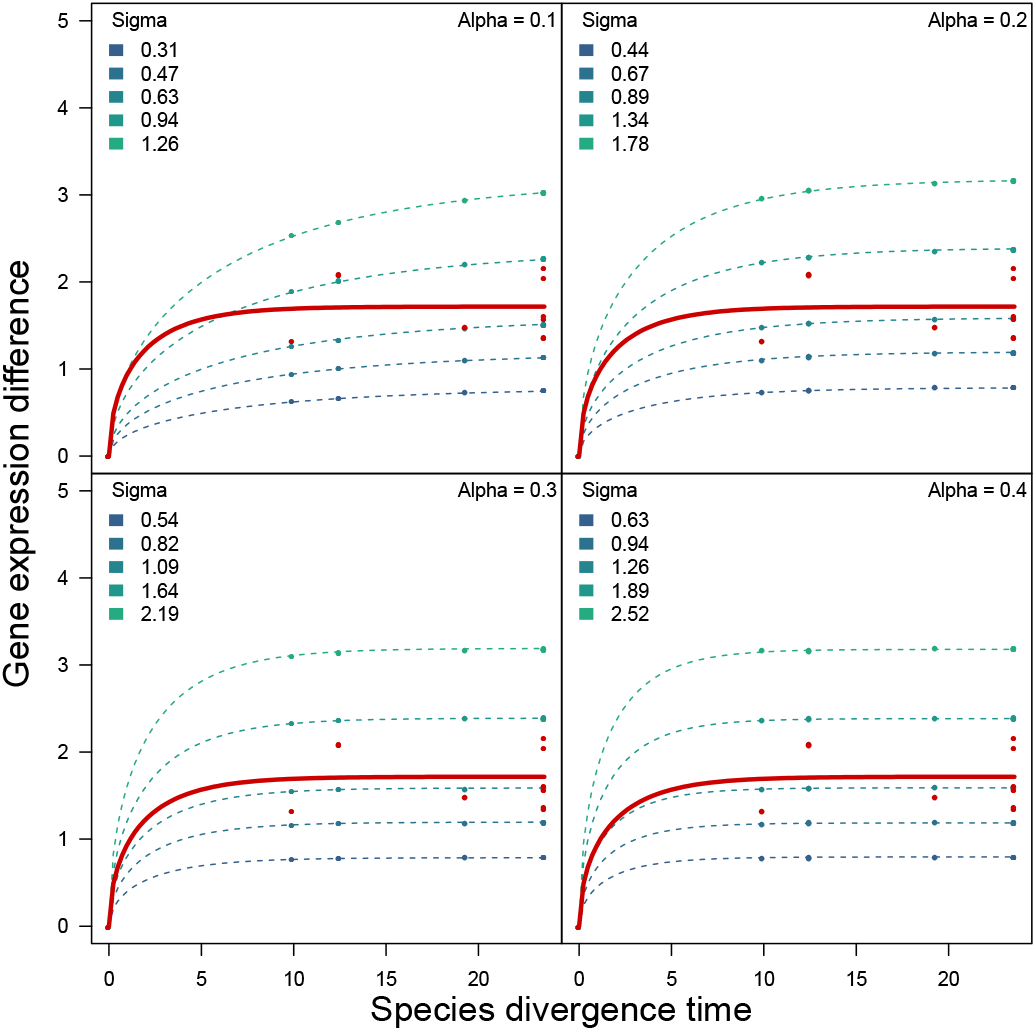
Between species gene expression variance plotted as a function of divergence time according to the *Heliconius* phylogeny. Red: σ^2^ from gene expression levels observed in *Heliconius*. Blue: simulated gene expression variance under different values of sigma. Each panel shows estimates for a different value of alpha (α).

### Gene expression evolution through genetic directional selection

To reveal branch-specific shifts in gene expression levels we applied BM and OU models, allowing for branch-specific shifts in the rest of the phylogeny. Using this approach, we found that 16% of the genes show a branch-specific shift, towards either up- or down-regulation, with increased expression levels showing lower variance than expected (Figure 8). The direction of gene expression shifts might be influenced by its mode of regulation. For example, in yeast it was found that regulatory mutations affecting *trans-*regulatory factors were more likely to cause an increase in gene expression. Conversely, mutations on *cis-*regulatory elements were found to be skewed towards a decrease in expression (Metzger et al. 2016). On the other hand, in primates, a higher proportion of species-specific gene expression shifts were found to be towards down-regulation (Khaitovich et al. 2005). If directional selection is causing a branch-specific shift in gene expression one would expect to see a low within-species variance, whereas if the shift is caused by a relaxation of purifying selection or perhaps balancing selection, a higher within-species variance would be expected.

When we looked at the degree of variability between genes showing a shift towards a higher or a lower expression level, we observed that down-regulated genes have a significantly higher variance than genes showing up-regulation (Figure 8). From this observation, we hypothesize that relaxation of purifying selection might be driving the shifts causing down-regulation on gene expression, a pattern which could eventually lead to a loss of expression. However, balancing selection or experimental noise could also lead to an elevated within-species variance. Because of the cost of gene expression, it is expected that only those genes that are essential and have fitness effects will continue to be expressed, whereas genes that are not will eventually stop being transcribed (Stern and Crandall 2018). However, a shift towards down-regulation does not always have to be a consequence of relaxed purifying selection. For example, in the orthocluster with id Group_449_clean_0, a 7-fold lower expression shift was detected in the branch leading to *H. doris* (Figure S5), and a significantly smaller variance than expected transcriptome-wide (Fisher’s exact test, *P*-value < 0.001). Directional selection favoring down-regulation of gene expression can occur in a scenario where fine tuning of expression levels are necessary for optimal cell or tissue function (Cayirlioglu et al. 2008; Catalán et al. 2016).

On the other hand, genes showing a branch-specific shift towards up-regulation have significantly lower variance when compared to expression level shifts towards down-regulation (Figure 8). This observed pattern could be a result of directional selection acting on gene expression levels leading to a reduction of the variation observed in gene expression. It is possible that in order to achieve an increase in gene expression levels, the selective forces leading to up-regulation would have to be sufficiently strong to result in a greater investment in energy allocated to transcription costs (Wagner 2005; Lang et al. 2009). Some of the genes having the most extreme branch shifts in expression, either toward a higher or a lower expression, are involved in enzymatic activity (Appendix I). Enzymes support biochemical and physiological processes helping in the optimization of tissue function (Wagner and Altenberg 1996; Feller and Gerday 1997). Thus, optimal enzymatic activity might be a key factor for species-specific brain and eye function, which in turn might be optimized for the species-specific life history and ecological environment.

A factor possibly influencing the proportion of transcriptome levels found to be evolving through drift, stabilizing or directional selection is the methodology used for orthology assessment. In our analysis of gene expression variation, we assessed variation in orthoclusters where an orthologous hit was found for each of our five *Heliconius* species. Genes with fast-evolving protein rates—to the point that orthology assessment becomes challenging—might also show gene expression shifts, which would not be detectable in our experimental design. For example, orthology assessment for genes showing sex-biased gene expression might require an alternative method. In fact, from the ortholclusters that we identified in this study, only two included genes with sex-biased expression (Catalán et al. 2018). Additionally, gene expression shifts due to duplication events need to be explored by applying an appropriate statistical approach, as expression of genes where a duplication event has happened could contribute to the fraction of the transcriptome evolving by directional selection.

With this work, we have generated a set of candidate genes that are putatively evolving through directional selection and that have the potential of being involved in the processes of adaptation and speciation. To test the role of these genes in such processes, functional validation will be necessary to gain a deeper insight in the evolutionary consequences of gene expression shifts. Techniques like *in situ* hybridization, RNAi and CRISPR/Cas9 are adequate tools that can be used in shedding light into the functionality of these genes. Particularly interesting could be those genes whose gene expression levels have shifted to the degree of showing absence of expression (Figure S6). The evolution of gain and loss of gene expression across a phylogeny requires a suitable theoretical framework that should be explored particularly since such events have the potential to accelerate evolution.

## Acknowledgements

Thanks to Aline Rangel Olguin for technical assistance with the orthocluster analysis. This work was partially supported by National Science Foundation grant IOS-1656260 to A.D.B and by the Knut & Alice Wallenberg Foundation to Jochen Wolf.

